# The *Drosophila* AWP1 ortholog Doctor No regulates JAK/STAT signaling for left-right asymmetry in the gut by promoting receptor endocytosis

**DOI:** 10.1101/2022.08.20.504629

**Authors:** Yi-Ting Lai, Sasamura Takeshi, Junpei Kuroda, Reo Maeda, Mitsutoshi Nakamura, Ryo Hatori, Tomoki Ishibashi, Kiichiro Taniguchi, Masashi Ooike, Tomohiro Taguchi, Naotaka Nakazawa, Shunya Hozumi, Takashi Okumura, Toshiro Aigaki, Mikiko Inaki, Kenji Matsuno

**Affiliations:** Department of Biological Sciences, Graduate School of Science, Osaka University, 1-1 Machikaneyama, Toyonaka, Osaka 560-0043, Japan; Department of Biological Science and Technology, Tokyo University of Science, 2641 Yamazaki, Noda, Chiba, 278-8510, Japan; Department of Biological Science, Tokyo Metropolitan University, 1-1 Minami-osawa, Hachioji, Tokyo 192-0397, Japan

**Keywords:** Left-right asymmetry, JAK/STAT signaling, *Drosophila* development, endocytic trafficking

## Abstract

Many internal *Drosophila* organs show stereotypical left-right (LR) asymmetry, for which the underlying mechanisms remain elusive. Here, we identified an evolutionarily conserved ubiquitin-binding protein, AWP1/Doctor no (Drn), as a novel factor required for the LR asymmetry of the embryonic anterior gut in *Drosophila*. We showed that *drn* is essential in the circular visceral muscle cells of the midgut for JAK/STAT signaling, which contributes to the first known cue for anterior gut lateralization via LR-asymmetric nuclear rearrangement. Embryos homozygous for *drn* and lacking its maternal contribution showed phenotypes similar to that of depleted JAK/STAT signaling, suggesting that Drn is a general component of JAK/STAT signaling. The absence of Drn resulted in the specific accumulation of Domeless (Dome), the receptor of JAK/STAT signaling, in intracellular compartments. Thus, Drn is required for the endocytic trafficking of Dome, which is subsequently degraded in lysosomes. Our results suggest that the endocytosis of Dome is a critical step in activating JAK/STAT signaling. The roles of AWP1/Drn in activating JAK/STAT signaling and in LR-asymmetric development may be conserved in various organisms.

**Summary Statement:** Dr. No, a *Drosophila* ortholog of AWP1, activates JAK/STAT signaling via Dome receptor endocytosis in a crucial step for left-right asymmetry in the developing gut.

## INTRODUCTION

Many animals show directional left-right (LR) asymmetry in body structure and function. Several mechanisms, such as cilia-generated flow, contribute to LR-axis formation in vertebrates (Yoshiba and Hamada, 2014). However, the mechanisms of LR-asymmetric development in invertebrates are relatively obscure and remain an elemental question in biology (Vandenberg and Levin, 2013).

Stereotypical LR asymmetry is seen in several organs in *Drosophila*, including the gut, testis, male genitalia, and brain (Coutelis et al., 2014; Okumura et al., 2008). Among these organs, the embryonic gut is the first structure to show LR asymmetry during development (Hayashi and Murakami, 2001; Hozumi et al., 2006). Intriguingly, the anterior and posterior parts of the embryonic gut are controlled by two distinct groups of genes. The LR asymmetry of the posterior part is defined by *Myosin31DF*, which determines cell chirality (Hozumi et al., 2006; Inaki et al., 2016; Inaki et al., 2018; Nakamura et al., 2013; Utsunomiya et al., 2019). An entirely different mechanism governs the LR asymmetry of the anterior part. The anterior gut of the embryo consists of the foregut (FG) and midgut (MG), which are complex structures with directional and stereotypical LR asymmetry (Fig. 1A, B) (Hayashi and Murakami, 2001). Various genetic pathways, including the JNK and Wnt pathways, play important roles in LR-asymmetric development (Kuroda et al., 2012; Ligoxygakis et al., 2001; Maeda et al., 2007; Okumura et al., 2010; Shin et al., 2021; Taniguchi et al., 2007). A genetic screen performed by our group suggested that the JAK/STAT signaling pathway is also involved in LR-asymmetric development. Our findings in the present study revealed the involvement of JAK/STAT signaling in the LR-asymmetric development of the anterior gut.

**Figure 1.**
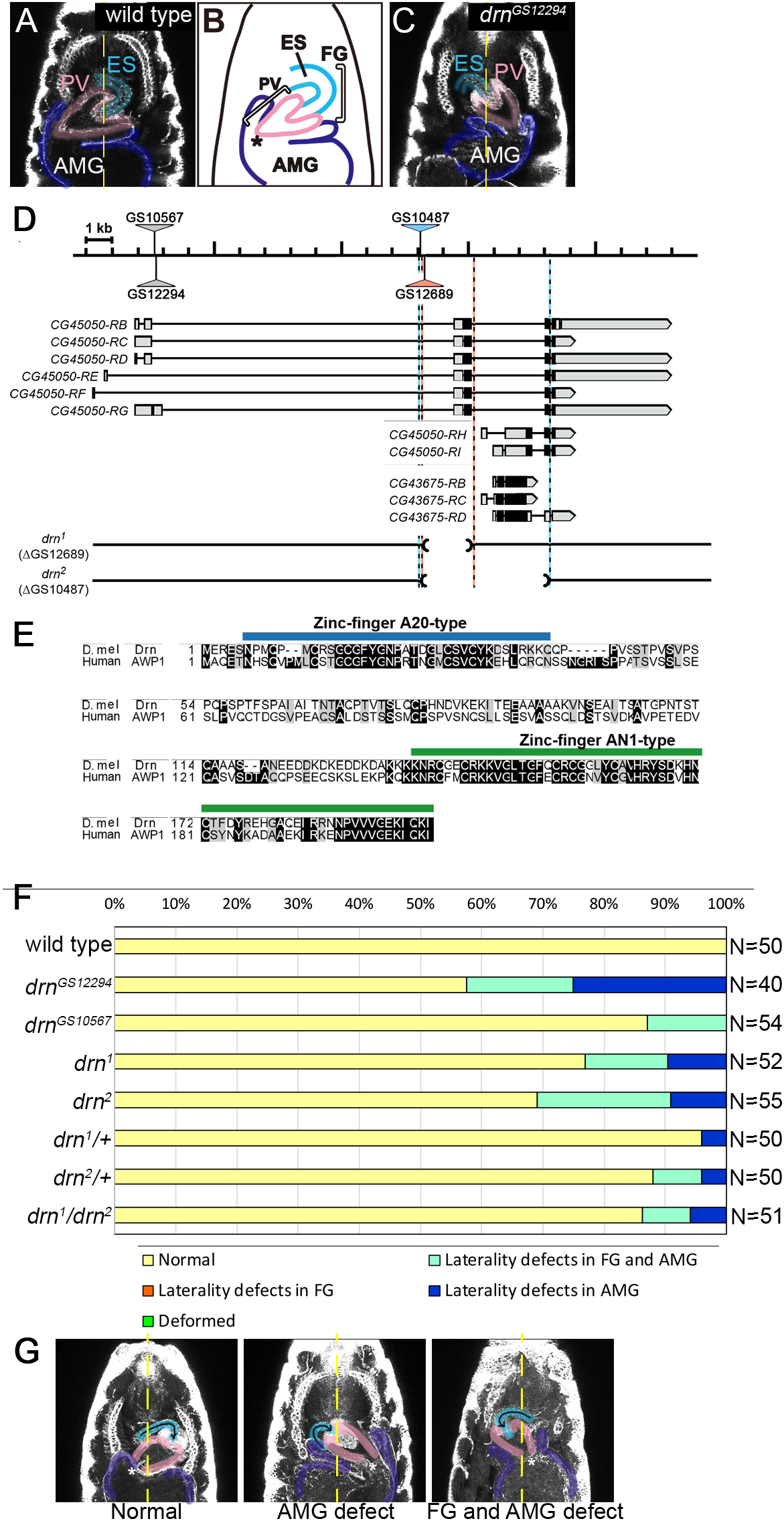
LR asymmetry of the FG and AMG is defective in *drn* mutants. (A) Ventral view of a stage-16 wild-type embryo. The gut outline was visualized by anti-Fas3 immunostaining. LR asymmetry in the FG was judged by the direction of ES (turquoise) rotation and in the AMG by the position of the joint (asterisk) between the AMG (dark blue) and the PV (pink) relative to the midline (yellow dotted line). (B) A diagram of the FG and AMG in wild-type embryos at stage 16, depicting the ES, PV, and AMG. (C) LR-inversion of the FG and AMG in *drn*^*GS12294*^ homozygotes. Symbols are the same as in A. (D) A diagram showing the genomic region of *drn* (upper), its alternative transcripts (middle), and the structures of *drn*^*1*^ and *drn*^*2*^ mutants (lower) with P-element insertion sites (triangles), deleted regions (parentheses and dotted lines), exons corresponding to coding regions (black boxes), untranslated regions (gray boxes), and introns (lines). (E) Amino acid sequences of *Drosophila* Drn (NP_001027162.1) and human AWP1 (NP_061879) were optimally aligned using ClustalW. The diagram indicates identical (white characters on black) and similar residues (black characters on gray). A20-type and AN1-type zinc finger domains are indicated by blue and green lines, respectively. (F) The frequency of FG and AMG LR defects (%) in embryos with the genotypes indicated at the left. Bars show the percentage of embryos with normal laterality (yellow); defects in laterality (reversed LR asymmetry or bilateral symmetry) in the AMG (blue) or in both the FG and AMG (turquoise), or deformities (green). The number (N) of embryos scored is shown at the right. (G) Examples of LR-asymmetry phenotypes in the FG and AMG in *drn*^*2*^ homozygotes. The ES (turquoise), PV (pink), and midline (yellow dotted line) are as described in A.

JAK/STAT signaling, which is evolutionarily conserved from *Drosophila* to humans, is essential in morphogenesis, cell proliferation, cell differentiation, cell death, immunity, and other biological events (Arbouzova and Zeidler, 2006; Calò et al., 2003; Lee et al., 2017; Myllymäki and Rämet, 2014; Recasens-Alvarez et al., 2017; Seif et al., 2017). In *Drosophila* JAK/STAT signaling, the ligands are encoded by *unpaired* (*upd*), *upd-2*, and *upd-3*, and the receptor is encoded by *domeless* (*dome*) (Agaisse et al., 2003; Brown et al., 2001; Harrison et al., 1998; Hombría et al., 2005). JAK is encoded by *hopscotch* (*hop*) in *Drosophila*, and as with the mammalian JAK/STAT system, Hop constitutively associates with Dome’s intracellular domain (Arbouzova and Zeidler, 2006; Binari and Perrimon, 1994). Ligand binding induces conformational changes in Dome that lead to the phosphorylation of JAK, which phosphorylates Dome to create docking sites for STATs—specifically, Stat92E in *Drosophila* (Yan et al., 1996). JAK then tyrosine-phosphorylates STATs, which are subsequently dimerized and translocated to the nucleus, where they bind enhancers of target genes and activate transcription (Arbouzova and Zeidler, 2006; Hou et al., 1996; Yan et al., 1996) (See Fig. 3A for diagram).

In mammals, endocytosis regulates JAK/STAT signaling through various mechanisms (German et al., 2011; Lei et al., 2011). Endocytosis and the regulation of JAK/STAT signaling activity are also closely connected in *Drosophila* (Devergne et al., 2007; Moore et al., 2020; Ren et al., 2015; Vidal et al., 2010). In the classic working model of endocytosis, membrane receptors are internalized into endosomal compartments where they are degraded and recycled, thereby reducing the number of receptors available to transduce signaling. (Cendrowski et al., 2016; Elkin et al., 2016; Piper et al., 2014). Aside from this, receptor activity requires endocytic trafficking in some signaling pathways (Cendrowski et al., 2016; Irannejad et al., 2013). Studies of the relationship between endocytosis and JAK/STAT signaling in *Drosophila* offer contradictory results as to whether endocytosis upregulates or downregulates JAK/STAT activity (Devergne et al., 2007; Moore et al., 2020; Ren et al., 2015; Vidal et al., 2010). To make sense of these conflicting results, it would be helpful to identify and examine a factor that specifically regulates the endocytosis of Dome.

Here, we found that the *Drosophila* ortholog of AWP1 (<u>a</u>ssociated <u>w</u>ith <u>P</u>RK<u>1</u>, also known as ZFAND6), referred to as Dr. No (Drn), positively regulates JAK/STAT signaling by facilitating the endocytic trafficking of the Dome receptor, which is required for the normal LR-asymmetric development of the embryonic gut. *Drosophila* Drn was named after Ian Fleming’s fictional character, whose heart was located in the right side of his chest (Fleming, 1958). Drn protein contains an A20-type zinc finger at the N-terminus and an AN1-type zinc finger at the C-terminus; this is similar to the mammalian ortholog, which was first identified in humans and mice (Duan et al., 2000). In vertebrates, AWP1 proteins bind ubiquitin, regulate NF-κB activity, and stimulate the export of Pex5 from the peroxisome, among other roles (Chang et al., 2011; Fenner et al., 2009; Miyata et al., 2012). During *Xenopus* development, AWP1 modifies Wnt and FGF signaling to specify neural crest cells (Seo et al., 2013). However, the molecular mechanisms by which AWP1/Drn proteins influence these various cell signaling pathways are not well understood. Here, we show that AWP1/Drn plays a crucial role in internalizing the Dome receptor, and propose a novel mechanism by which AWP1/Drn positively modulates JAK/STAT signaling.

## RESULTS

### *drn* mutations affect LR-asymmetric gut morphogenesis in *Drosophila*

To identify genes that affect LR asymmetry in the anterior gut, including the FG and anterior midgut (AMG), we conducted a genetic screen using a large collection of P-element insertion lines (*Drosophila* Genes Search Project, http://kyotofly.kit.jp/stocks/documents/GS_lines.html) (Toba et al., 1998). We scored the LR-asymmetry phenotypes of the anterior gut in mutants obtained from our genetic screen. The FG is composed of the pharynx, the esophagus (ES), and the proventriculus (PV), which is a valve-like structure connecting the FG to the AMG (Fig. 1A, B). We defined normal LR asymmetry in the FG and AMG as follows: 1) when viewed from the ventral side, the wild-type *Drosophila* ES loops in an inverse C shape and is connected to the PV (100%, N=50); this was defined as normal FG laterality (Fig. 1A). 2) the joint between the PV and AMG is located on the right side of the midline in wild-type embryos (100%, N=50); this was defined as normal laterality of the AMG (Fig. 1A). Using these criteria, our genetic screen identified two mutant lines, GS12294 and GS10567, that affect LR asymmetry in the FG and AMG (Fig. 1C).

These lines carry a P-element insertion in the *CG45050* locus (Fig. 1D). In this study, we named the *CG45050* gene *dr. no* (*drn*). *Drosophila* Drn and human AWP1 share 42.1% identity and 62.2% similarity for the whole protein (calculated with EMBOSS Pairwise Alignment Algorithms) (Fig. 1E). In particular, an A20-type zinc finger (amino acids 6-40) and an AN1-type zinc finger (amino acids 137-180) are highly conserved (Fig. 1E). A20-type zinc fingers are found in various proteins with ubiquitin-editing functions; these proteins are often associated with human pathogenesis (Heyninck and Beyaert, 1999; Jacque and Ley, 2009; Kim et al., 2021; Rothe et al., 1995; Song et al., 1996). We found LR-asymmetry defects of the anterior gut in 42.5% of *Drosophila* embryos homozygous for *drn*^*GS12294*^ and 13.0% of those homozygous for *drn*^*GS10567*^, indicating a disturbance in LR-asymmetrical development (Fig. 1C, F). In contrast, hindgut and posterior midgut laterality were normal in all cases examined (N=40), indicating heterotaxy but not situs inversus in these phenotypes. To investigate Drn function, we generated null mutant alleles of *drn* by imprecise P-element excision. In *drn*^*1*^, the deduced initiation codon and 5’ portion of the coding region were deleted from the *drn* alternative RNA products CG45050-RB, -RC, -RD, -RE, -RF, and -RG (Fig. 1D). In *drn*^*2*^, the deduced initiation codon and most of the coding sequence were deleted from all of the *drn* alternative RNA products, suggesting that *drn*^*2*^ is a null mutation of *drn* (Fig. 1D). Various degrees of LR-asymmetry defects were observed in *drn*^*1*^ or *drn*^*2*^ homozygotes and trans-heterozygotes, demonstrating that mutations in *drn* are responsible for the LR-defect phenotypes (Fig. 1F, G). These mutant embryos had LR defects in both the FG and AMG or in the AMG alone, but never in the FG alone (Fig. 1F, G). Thus, LR-asymmetric defects in the FG are strictly coupled with those in the AMG, suggesting the primary role of the AMG in the LR-asymmetric development of the anterior gut as has been observed in other mutants with defective LR asymmetry (Kuroda et al., 2012; Taniguchi et al., 2007). We also found relatively mild LR defects in heterozygotes of *drn*^*1*^ or *drn*^*2*^, indicating that *drn* might behave in a semi-dominant manner (Fig. 1F).

### *drn* is required for the LR-asymmetric rearrangement of CVMU cells

To confirm that *drn* is required in the LR-asymmetric development of the anterior gut, we performed rescue experiments using the GAL4/UAS system (Brand and Perrimon, 1993). Expression of a wild-type *drn* encoded by *UAS-drn* was driven by various tissue-specific GAL4 drivers in a *drn*^*1*^ homozygote mutant background (Brand and Perrimon, 1993; Elliott and Brand, 2008). The FG and AMG are composed of the epithelium (Epi), circular visceral muscle (CVMU), and longitudinal visceral muscle (LVMU) (Fig. 2A). LR defects of the FG and AMG in *drn*^*1*^ homozygotes were reduced by half when *UAS-drn* was introduced without any GAL4 driver (negative control), which is probably due to leaky expression of *UAS-drn* (Fig. 1F, 2B). On the other hand, these LR defects were efficiently rescued by *UAS-drn* misexpression driven by *da*-GAL4 (ubiquitous), *NP1522* (in the somatic muscle and CVMU), *hand*-GAL4 (in the CVMU), *24B*-GAL4 (in the CVMU, LVMU, and somatic muscle), and *48Y*-GAL4 (in the AMG epithelium, the CVMU, and the LVMU) (Fig. 2B). In contrast, LR defects were not rescued by *UAS-drn* misexpression driven by *NP0221* (in the LVMU), *NP5021* (in the epithelium), or *elav-GAL4* (in the nervous system) (Fig. 2B). Collectively, our data suggest that *drn* expression is required primarily in the CVMU, not in the LVMU or other tissues, for normal LR-asymmetric development of the FG and AMG. We also found that in wild-type embryos, *UAS-drn* misexpression driven by the GAL4 drivers tested here did not induce marked LR defects in the FG and AMG, suggesting that enhanced *drn* expression did not affect LR-asymmetric development in wild-type embryos (Fig. S1).

**Fig. 2.**
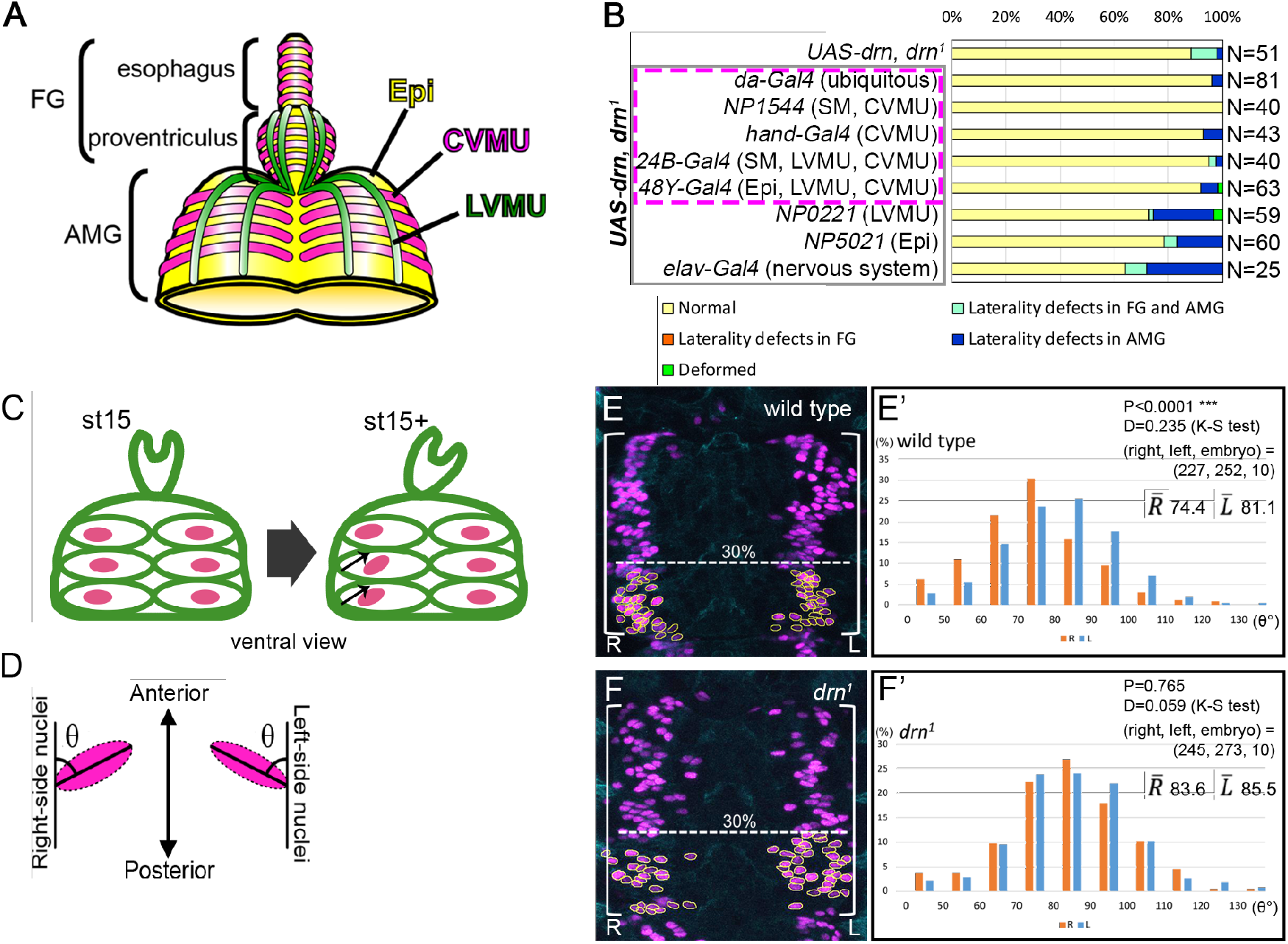
*drn* is required in the CVMU of the midgut for the LR-asymmetric development of the FG and AMG. (A) A diagram of the visceral muscles after stage 14: the FG and AMG are overlaid by a layer of CVMU (magenta), and the PV and AMG are covered by LVMU (green). The epithelium is shown in yellow. (B) The frequency of LR-asymmetry defects (%) in the FG and AMG in embryos carrying *UAS-drn, drn*^*1*^/*drn*^*1*^ without (control, *UAS-drn, drn*^*1*^) or with the GAL4 drivers outlined in gray at the left; *GAL4* expression was driven in the cell types or tissues shown in parentheses (SM: somatic muscles; Epi: epithelium of the AMG). Bars show the percentage of embryos with no laterality defects (yellow), laterality defects in the AMG (blue) or both the FG and AMG (turquoise), or with deformities (green). *UAS-drn* misexpression driven by the GAL4 drivers outlined in magenta rescued the laterality defects. The number (N) of embryos scored is shown at the right. (C) A diagram showing the LR-asymmetric rearrangement of nuclei (magenta) in CVMU cells (green ellipses) of the AMG from early (st15) to late stage 15 (st15+). Small arrows indicate the tilt of the ellipsoid nuclei. (D) A diagram showing the major axis angles of the ellipsoid nuclei, represented by the angle (θ) between the major axis of the right- and left-side ellipsoid nuclei and the anterior-posterior axis of the embryo. (E and F) Ventral views of the AMG showing CVMU cells and their nuclei, stained by anti-Fas3 (cyan) and anti-RFP (magenta) antibodies, respectively, in control (E, *65E04-GAL4*/*UAS-Redstinger*) and *drn*^*1*^ homozygote (F, *drn*^*1*^, *65E04-GAL4*/*drn*^*1*^, *UAS-Redstinger*) embryos at late stage 15. Nuclei in the lower 30% of the presumptive first chamber (indicated by white angle brackets) were selected for measurement (encircled by yellow lines). (E’, F’) Frequency histograms of the axis angles (in 10-degree increments) of the left (blue bars) and right (orange bars) sides of CVMU cell nuclei in the ventral AMG of control (E’, *65E04-GAL4*/*UAS-Redstinger*) and *drn*^*1*^ homozygote (F’, *drn*^*1*^, *65E04-GAL4*/*drn*^*1*^, *UAS-Redstinger*) embryos at late stage 15. P-values (upper-right corner) indicate the statistical significance of differences between the angle distributions of the right and left sides calculated by K-S test. Numbers in parentheses indicate the numbers (right nuclei, left nuclei, and embryos) analyzed. Average angles of the right and left sides are indicated as 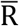 and 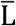, respectively.

We previously showed that the LR-asymmetric tilt of the ellipsoidal nuclei in the CVMU cells covering the midgut epithelium is the first point of disruption of LR symmetry in the anterior gut (Fig. 2C) (Kuroda et al., 2012; Okumura et al., 2010; Shin et al., 2021; Taniguchi et al., 2007). At early-stage 15, the major axis of the ellipsoidal nuclei in the CVMU cells is still perpendicular to the midline of the AMG (Fig. 2C). When the midgut chambers began dividing at late stage 15, the nuclei in the lower 30% of the presumptive first chamber on the right began tilting diagonally up and to the right (Fig. 2C). This LR-asymmetric nuclear tilt precedes the LR-asymmetric morphological changes of the AMG, indicating that it is not a consequence of LR-asymmetric morphogenesis (Taniguchi et al., 2007). To analyze whether this process was disrupted in *drn* mutants, we used *65E04-GAL4* to specifically express nuclear-localizing RedStinger fluorescent proteins in the visceral muscles. The angle (θ) between the major axis of the ellipsoidal nuclei in the lower 30% of the presumptive first chamber and the embryo’s anterior-posterior axis was measured by double-blind test at late stage 15 (Fig. 2D). As with our previous results, the angle measured was smaller for nuclei in the right side than those in the left in wild-type embryos (*P*<0.0001, K-S test) (Fig. 2E, E’) (Kuroda et al., 2012; Okumura et al., 2010; Taniguchi et al., 2007). However, in *drn*^*1*^ homozygotes, there was no significant difference in angle between the left and right sides, and the angle remained closer to perpendicular on both sides even at later stage 15 (P=0.765, K-S test) (Fig. 2F, F’). In addition to the LR-asymmetric tilting of the nuclei, we previously reported that the nuclei assemble in belt-shaped zones along the anterior-posterior axis. These nuclei are scattered in mutants with defects in the LR asymmetry of the anterior gut (Shin et al., 2021). We here observed that the distribution of the nuclei was more dispersed in *drn*^*1*^ homozygotes than in wild-type embryos in all cases examined (N=10) (Fig. 2E, F). Collectively, these results suggest that *drn* contributes to the LR-asymmetric development of the AMG by regulating the LR-asymmetric rearrangement of nuclei in CVMU cells.

To verify the roles of *drn* in the CVMU, we analyzed the distribution of *drn* mRNA in embryos by *in situ* hybridization at various stages of embryogenesis (Fig. S2A-D). Comprehensive analyses of gene expression in *Drosophila* show that *drn* is highly expressed in early embryonic stages (Thurmond et al., 2019). We found that *drn* mRNA was strongly detected in pre-blastoderm to blastoderm (stage 5) stages, suggesting that *drn* mRNA is maternally provided (Fig. S2A, B). We also found that *drn* was broadly expressed at stages 11 and 15, including the trunk visceral mesoderm (TVM, the primordium of the CVMU), where *drn* was required for the LR-asymmetric development of the anterior gut (Fig. S2C). In contrast, a negative control (sense probe) gave no signal under the same conditions (Fig. S2E-H).

### JAK/STAT signaling is involved in the LR-asymmetric development of the anterior gut

Although previous research suggested that Drn negatively regulates JAK/STAT signaling in cultured *Drosophila* cells (Vidal et al., 2010), the role of Drn *in vivo* was not further explored. Considering that *drn* is involved in JAK/STAT signaling, we hypothesized that JAK/STAT signaling plays a role in the LR-asymmetric morphogenesis of the anterior gut. Various mutants of genes involved in the JAK/STAT signaling pathway, including *dome, Stat92E, upd*, and *hop*, were scored for LR phenotypes (Fig. 3B). The roles of these gene products are schematically shown in Fig. 3A. These mutants showed various degrees of LR defects in the FG and AMG, indicating that JAK/STAT signaling is indispensable for normal LR-asymmetric development of the anterior gut (Fig. 3B). In addition, when JAK/STAT signaling was augmented by misexpressing an activated form of Hop (*UAS-hop*^*Tum-l*^) (Harrison et al., 1995) specifically in CVMU cells under the control of *hand-GAL4* or *24B-GAL4*, the embryos also showed LR defects in the FG and AMG (Fig. 3B). Therefore, excessive activation of JAK/STAT signaling also disrupts the LR asymmetry of the FG and AMG. Taken together, these findings indicate that JAK/STAT signaling activity must be maintained at proper levels for the LR-asymmetric development of the anterior gut.

**Fig. 3.**
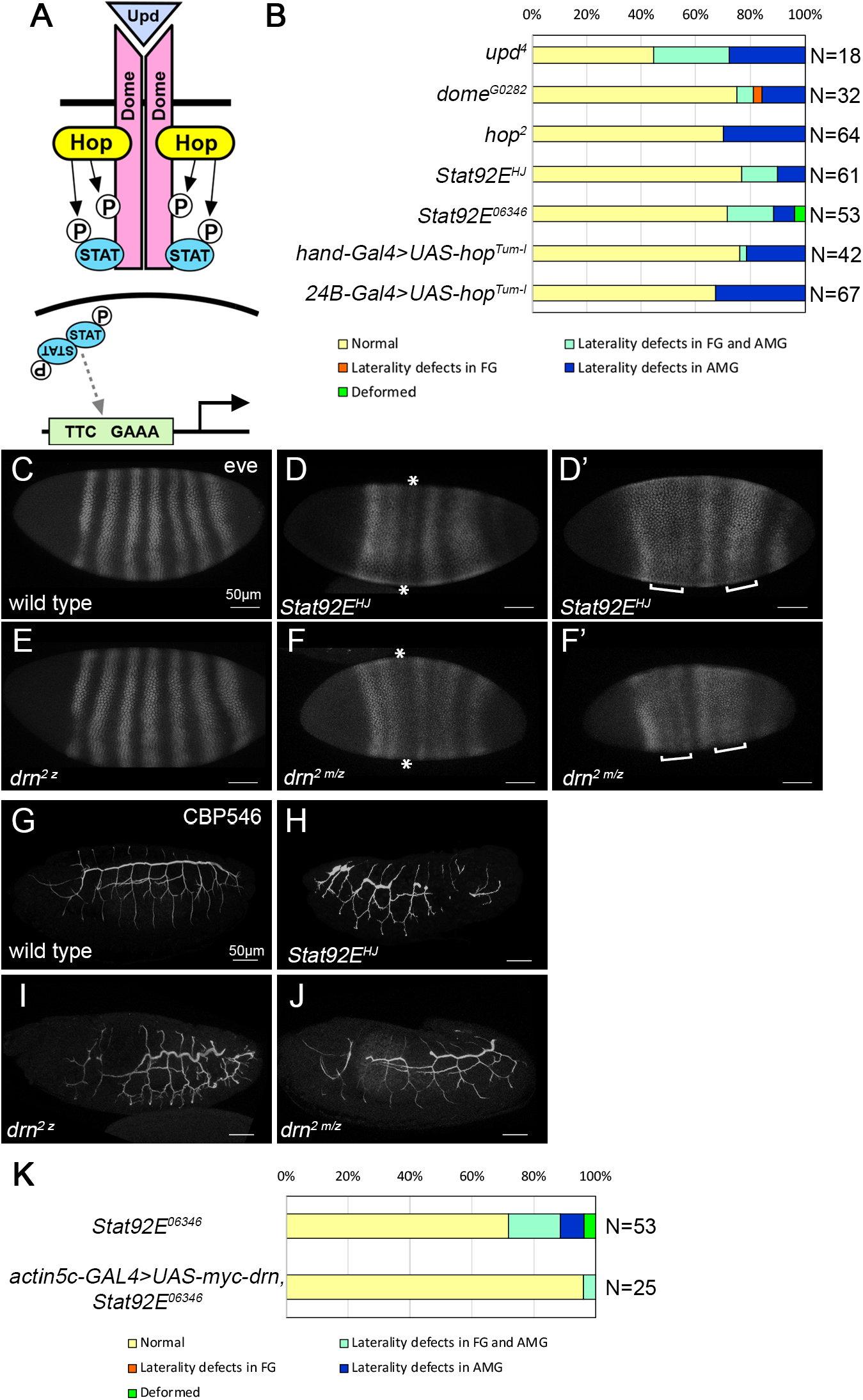
*drn* functions collectively with JAK/STAT signaling during embryonic development. (A) Illustration of the *Drosophila* JAK/STAT pathway. (B) Frequency of LR defects (%) in the AMG of embryos with the genotypes indicated at the left. Bars show the percentage of embryos with normal laterality (yellow), laterality defects in the FG and AMG (turquoise), laterality defects only in the AMG (blue), and deformities (green), respectively. The number (N) of embryos scored is shown at the right. (C-F’) The expression of *even-skipped* in wild-type (C), *Stat92E*^*HJ*^ (D and D’), *drn*^*2 z*^ (zygotic mutant) (E), and *drn*^*2 m/z*^ (zygotic and maternal mutant) (F and F’) embryos. D and F show the phenotype of weakened stripe 3 (marked with asterisks). D’ and F’ show the phenotype of stripe fusion (marked with brackets). (G-J) The trachea was detected by CBP546 staining in wild-type (G), *Stat92E*^*HJ*^ (H), *drn*^*2 z*^ (zygotic mutant) (I), and *drn*^*2 m/z*^ (zygotic and maternal mutant) (J) embryos. (K) The frequency of LR-asymmetry defects (%) observed in the FG and AMG of *Stat92E*^*06346*^ homozygotes without (control, upper) and with ubiquitous misexpression of *UAS-myc-drn* driven by *actin5c-GAL4* (lower). Bars show the percentage of embryos with normal laterality (yellow), laterality defects in the FG and AMG (turquoise), laterality defects in only the AMG (blue), and deformities (green), respectively. The number (N) of embryos scored is shown at the right. Scale bars in C-J: 50 μm.

### *drn* is a general component of the JAK/STAT signaling pathway

To analyze the connection between *drn* and JAK/STAT signaling more directly, we assessed the characteristic phenotypes relating to JAK/STAT signaling in *drn* mutants. The expression of a pair-rule gene, *even-skipped* (*eve*), is disturbed in embryos homozygous for *Stat92E*, such as *Stat92E*^*HJ*^, with about 67% of the embryos showing aberrant phenotypes such as weak stripe 3 (19%) or fusion between stripe 2 and 3 (19%) as well as fusion between stripe 5 and 6 (67%) (N=27), rather than the seven-stripe pattern seen in wild-type embryos (Fig. 3C-D’) (Yan et al., 1996). We next analyzed *eve* expression in embryos homozygous for *drn*^*2*^, and found that al embryos tested had normal *eve* expression and displayed a normal seven-stripe pattern (N=20) (Fig. 3E). Since we found that the mRNA of *drn* is maternally supplied (Fig. S2A, B), we genetically removed maternal *drn* from *drn*^*2*^ homozygotes (*drn*^*2 m/z*^). The phenotypes of *drn*^*2 m/z*^ embryos were similar to those of *Stat92E*^*HJ*^ homozygotes (36%, N=33) (Fig. 3F, F’). These results suggest that *drn* is required for JAK/STAT signaling where Stat92E plays an essential role. To further verify this idea, we examined trachea morphology, which is controlled by JAK/STAT signaling in later embryonic stages (st 15-17) (Li et al., 2003). The trachea was detected by CBP546 staining (Dong et al., 2014). In *Stat92E*^*HJ*^ homozygotes, the dorsal trunk of the trachea was disturbed as compared to that of wild-type embryos (Fig. 3G, H) and was truncated in some *drn*^*2*^ homozygotes (35%, N=20) and *drn*^*2 m/z*^ embryos (33%, N=21). Defects in *drn*^*2 m/z*^ embryos were more severe than in *drn*^*2 z*^ homozygotes, which can be predicted from the maternal contribution of *drn* (Fig. 3I, J). These results suggested that as with *Stat92E, drn* plays a positive role in JAK/STAT signaling during embryonic development.

To further verify this possibility, we assessed whether the ubiquitous misexpression of *drn*, driven by *actin5c-GAL4*, could rescue LR defects in *Stat92E*^*06346*^ homozygous embryos. Embryos homozygous for *Stat92E*^*06346*^ showed LR defects in the anterior gut at a frequency of 25% (Fig. 3K); however, ubiquitous misexpression of *drn* reduced the frequency of LR defects to 4% (Fig. 3K). This result was not due to epistasis between *drn* and *Stat92E*, since embryos homozygous for *Stat92E*^*06346*^ still have residual *Stat92E* activity, which is maternally supplied (Hou et al., 1996; Li et al., 2003; Tsurumi et al., 2011). Nevertheless, our studies demonstrated that *drn* positively contributes to activate JAK/STAT signaling in three different contexts of development. Therefore, we here propose that Drn is a general component positively acting on JAK/STAT signaling, although a previous study involving knockdown by RNA interference (RNAi) and a reporter assay showed that wild-type *drn* downregulated JAK/STAT signaling in *Drosophila* cultured cells (Vidal et al., 2010). The cause of this discrepancy is unclear, but may involve a negative feedback loop that is active in a particular time frame in JAK/STAT signaling detected by the reporter assay in cultured cells.

### Drn may interact with Rab GTPases in the endocytic trafficking pathway

Although the biochemical roles of Drn have not been studied in *Drosophila*, the mammalian ortholog AWP1 binds ubiquitin and modulates functions of ubiquitinated proteins in mammals (Chang et al., 2011; Duan et al., 2000; Fenner et al., 2009); this process is often related to endocytic trafficking and the lysosomal breakdown of membrane receptors (Piper et al., 2014). Thus, we hypothesized that Drn might be involved in endocytic trafficking that regulates the JAK/STAT signaling pathway. Since TVM and CVMU cells are located deep inside the embryo, it is difficult to obtain clear microscopic images, making them unsuitable for analyzing Drn’s subcellular localization. However, *in situ* hybridization analysis revealed that *drn* is also expressed in the embryo’s epidermis (Fig. S2). Thus, we analyzed the potential colocalization of Drn with various endocytic compartments in the epidermis.

To detect Drn protein, we generated a polyclonal antibody (anti-Drn antibody) against a full-length Drn and assessed its specificity using the UAS-GAL4 system and RNAi against *drn* to deplete Drn protein in the stripe along the anterior-posterior boundary of the wing disc (the region expressing *ptc*). We found that anti-Drn antibody staining was largely absent from the stripe, confirming the antibody’s specificity (Fig. S3). Using this anti-Drn antibody, we examined the potential colocalization of Drn with various endosomal compartments in the epidermis of wild-type embryos. Drn was detected as punctae in the cytosol and was occasionally found colocalized with endosomal markers such as Hrs (early endosomes), Rab5 (early endosomes), Rab7 (late endosomes), LAMP1 (lysosomes), and Rab11 (recycling endosomes) (white arrowheads in Fig. 4A-E’’). Thus, although Drn distribution did not concentrate with any particular endosomal markers, Drn appeared to localize to endocytic compartments such as Hrs, Rab5, Rab7, and LAMP1 at a low frequency, which is consistent with our idea that Drn has some role in endocytic trafficking.

**Fig. 4.**
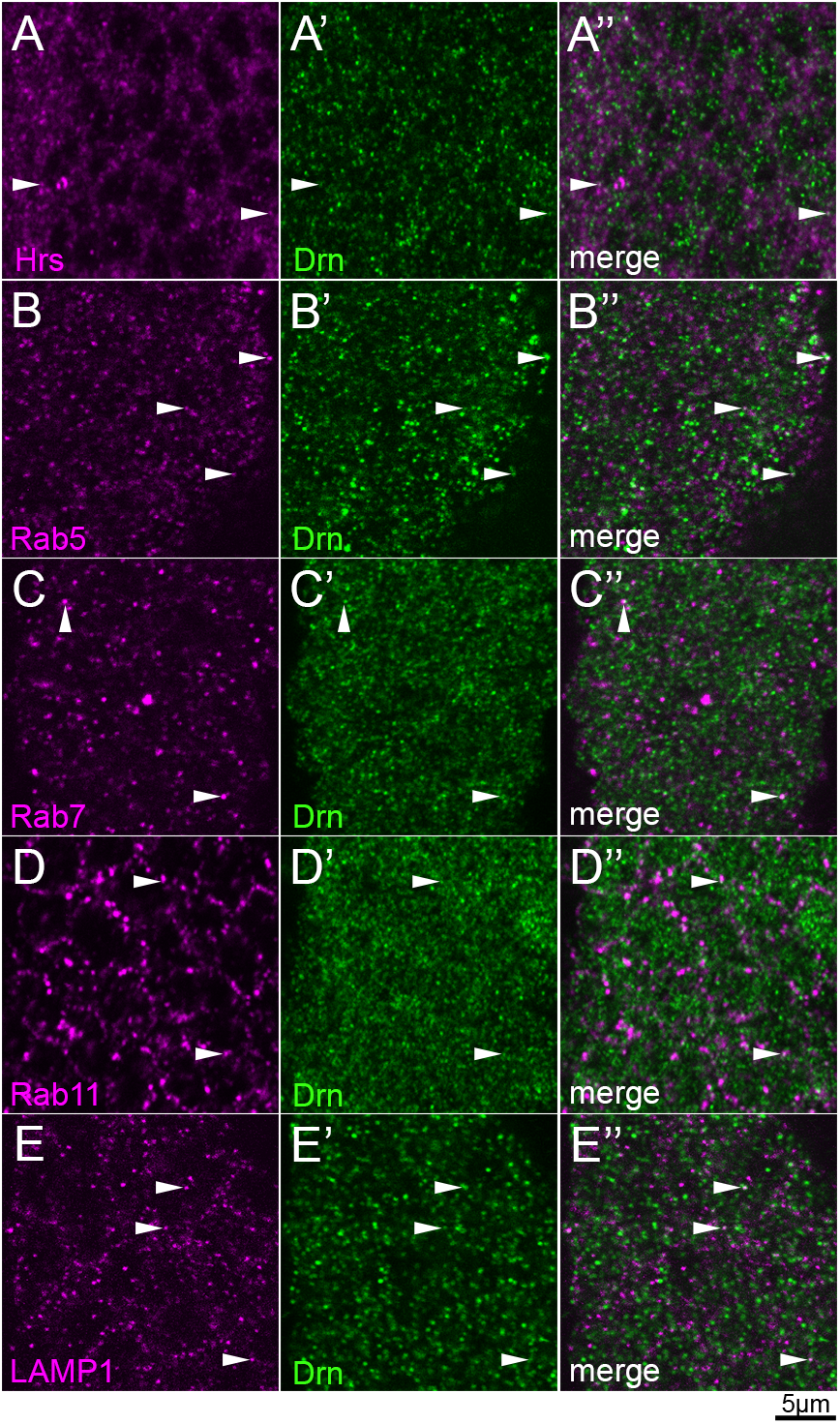
Drn occasionally colocalized with markers of various endocytic compartments in the epidermis of wild-type embryos. (A-E”) Drn subcellular localization in the epidermal cells of wild-type embryos. Embryos were double-stained with anti-Drn (green, middle and right columns) and markers for the following endosome markers (magenta, left and right columns): (A, and A”) Hrs (early endosomes); (B and B”) Rab5 (early endosomes); (C and C”) Rab7 (late endosomes); (D and D”) Rab11 (recycling endosomes); (E and E”) LAMP1 (lysosomes). (A”-E”) show merged images of A-E and A’-E’, respectively. White arrowheads indicate vesicles showing the colocalization of Drn with markers of various endocytic compartment. Scale bar: 5 μm.

### Drn is required for endocytic trafficking of the Dome receptor

Given that Drn positively contributes to JAK/STAT signaling by regulating endocytic trafficking, we speculated that Drn might regulate the JAK/STAT signaling receptor Dome during its endocytic trafficking (Brown et al., 2001). We analyzed the subcellular distribution of Dome using a full-length Dome protein that has a C-terminal GFP tag (Dome-GFP) and maintains wild-type Dome functions (Ghiglione et al., 2002). *UAS-dome-GFP* was ubiquitously driven under the control of *da-GAL4* in wild-type and *drn* homozygous embryos (Fig. 5A-B”).

**Fig. 5.**
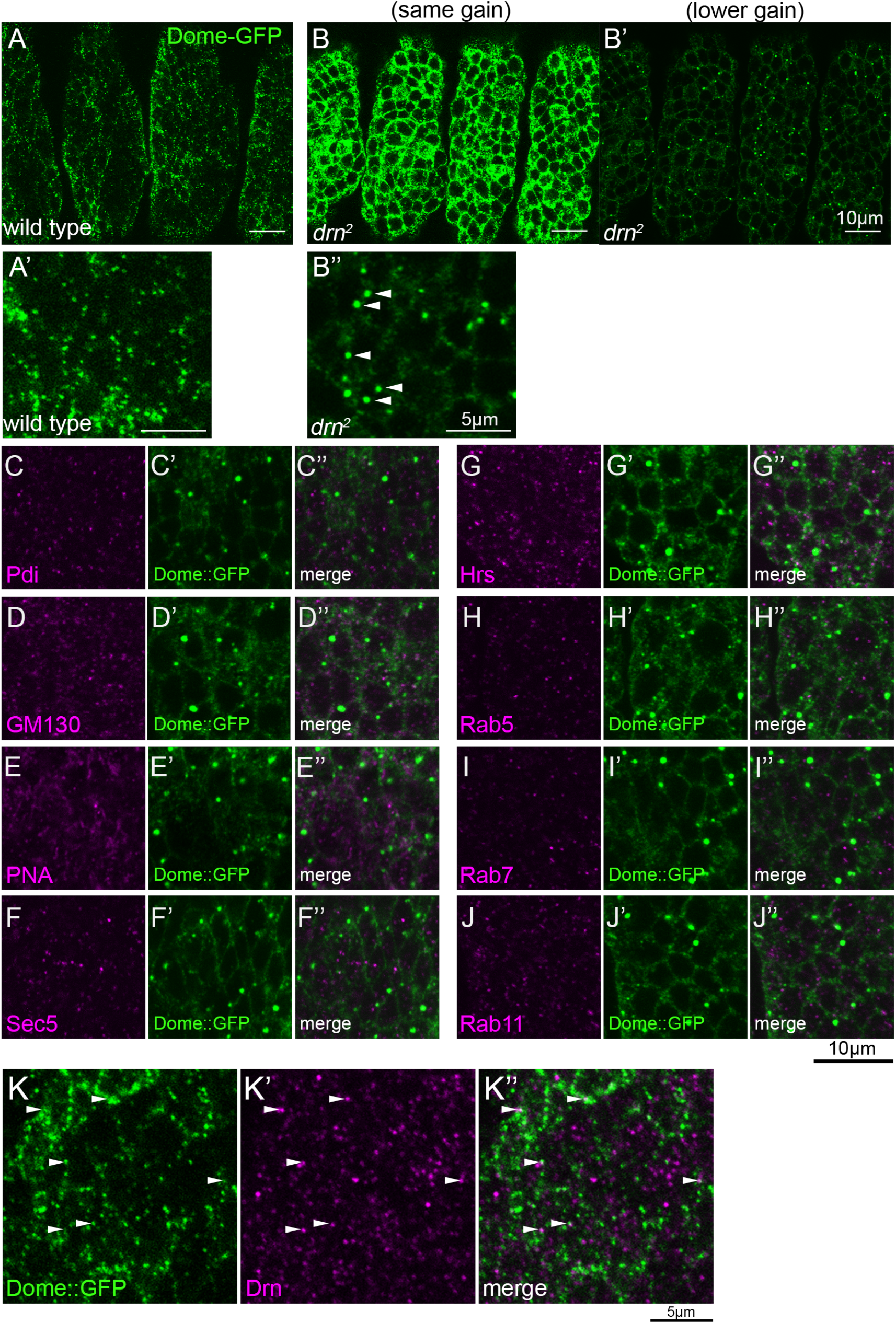
Dome accumulated in the cell cortex and intracellular structures in the epidermis of *drn* mutant embryos. (A-B’’) Ubiquitous *UAS-dome-GFP* expression was driven by *da-GAL4*. Dome-GFP was detected by anti-GFP antibody staining in the epidermal cells of wild-type (A) and *drn*^*2*^ (B-B’’) embryos. Gain to capture was the same for images in A and B; images in B’ and B” use lower gain to visualize intracellular aggregations and avoid signal saturation. A’ and B” are magnified views of A and B’. White arrowheads in B” indicate Dome-GFP aggregations. Scale bars: 10 μm in A-B’; 5 μm in A’ and B”. (C-J”) The subcellular localization of Dome-GFP aggregates in epidermal cells of *drn*^*2*^ mutants. Embryos were double-stained with anti-GFP antibody (green, middle and right columns) and antibodies against the following markers of intracellular compartments (magenta, left and right columns): (C, C”) Pdi (ER); (D, D”) GM130 (*cis*-Golgi); (E, E”) PNA (*trans*-Golgi); (F, F”) Sec5 (exocyst); (G, G”) Hrs (early endosome); (H, H”) Rab5 (early endosome); (I, I”) Rab7 (late endosome); (J, J”) Rab11 (recycling endosome). C”-J” show merged images of C-J and C’-J’, respectively. Scale bar: 10 μm. (K-K’’) Expression of *UAS-dome-GFP* was driven ubiquitously by *da-GAL4* in wild-type, and Dome-GFP and Drn were detected by anti-GFP (green, K and K”) and anti-Drn (magenta, K’ and K”) antibody staining in epidermal cells. K” is a merged image of K and K’. White arrowheads indicate vesicles showing colocalization between Dome-GFP and Drn. Scale bar: 5 μm.

In wild-type embryos, Dome-GFP was observed in epidermal cells as punctae localized to the vicinity of the plasma membrane and to cytosolic vesicles (Fig. 5A, A’; Fig. S4). Cytoplasmic vesicles containing Dome-GFP were occasionally labeled by markers for intracellular components, such as Rab5 and Rab11, but Dome did not appear to associate with any particular intracellular components (Fig. S4). In all cases, Dome-GFP was markedly increased in the epidermal cells of *drn*^*2*^ homozygotes compared to wild-type embryos, as observed in images of Dome-GFP obtained at the same gain of signal (Fig. 5B). In images obtained by reduced gain of signal detection, Dome-GFP was observed as larger clumps located near the plasma membrane (Fig. 5B’, B”). To analyze the nature of these clumps, we co-stained Dome-GFP with markers for various intracellular compartments, including Pdi (ER), GM130 (*cis*-Golgi), PNA (*trans*-Golgi), Sec5 (exocytic vesicle), Hrs (early endosomes), Rab5 (early endosomes), Rab7 (late endosomes), and Rab11 (recycling endosomes) in the epidermal cells of *drn*^*2*^ homozygotes. None of these markers colocalized with Dome-GFP (Fig. 5C-J”). Nevertheless, the large aggregations of Dome-GFP in *drn* mutants suggests a failure of Dome endocytic trafficking, which may prevent the degradation of Dome-GFP in lysosomes. Such a defect in endocytosis may cause JAK/STAT signaling to deteriorate, supporting previous suggestions that endocytosis is essential for Dome activation (Devergne et al., 2007; Moore et al., 2020). Unlike Dome-GFP, the Wnt and Notch signaling receptors Frizzled2 and Notch, respectively, were unchanged in the epidermis of wild-type and *drn*^*2*^ homozygotes (Fig. S5). Hence, the defective endocytosis in *drn*^*2*^ homozygotes is specific to Dome-GFP.

Considering that Drn may facilitate the endocytic trafficking of Dome, we considered whether Dome-GFP and Drn might colocalize in some intracellular vesicles. Although the staining patterns of Drn and Dome-GFP did not broadly resemble each other, they colocalized in a small population of vesicles (white arrowheads in Fig. 5K-K”). These results suggest that Drn may directly or indirectly interact with Dome in some endocytic compartments to facilitate proper Dome trafficking. This agrees with the finding that Dome is ubiquitinated (Tognon et al., 2014). Taking these results together, we speculate that, as seen in other organisms, the binding of ubiquitin to Drn specifically promotes the internalization of Dome, which is a critical step for activating JAK/STAT signaling (Fig. 6).

**Fig. 6.**
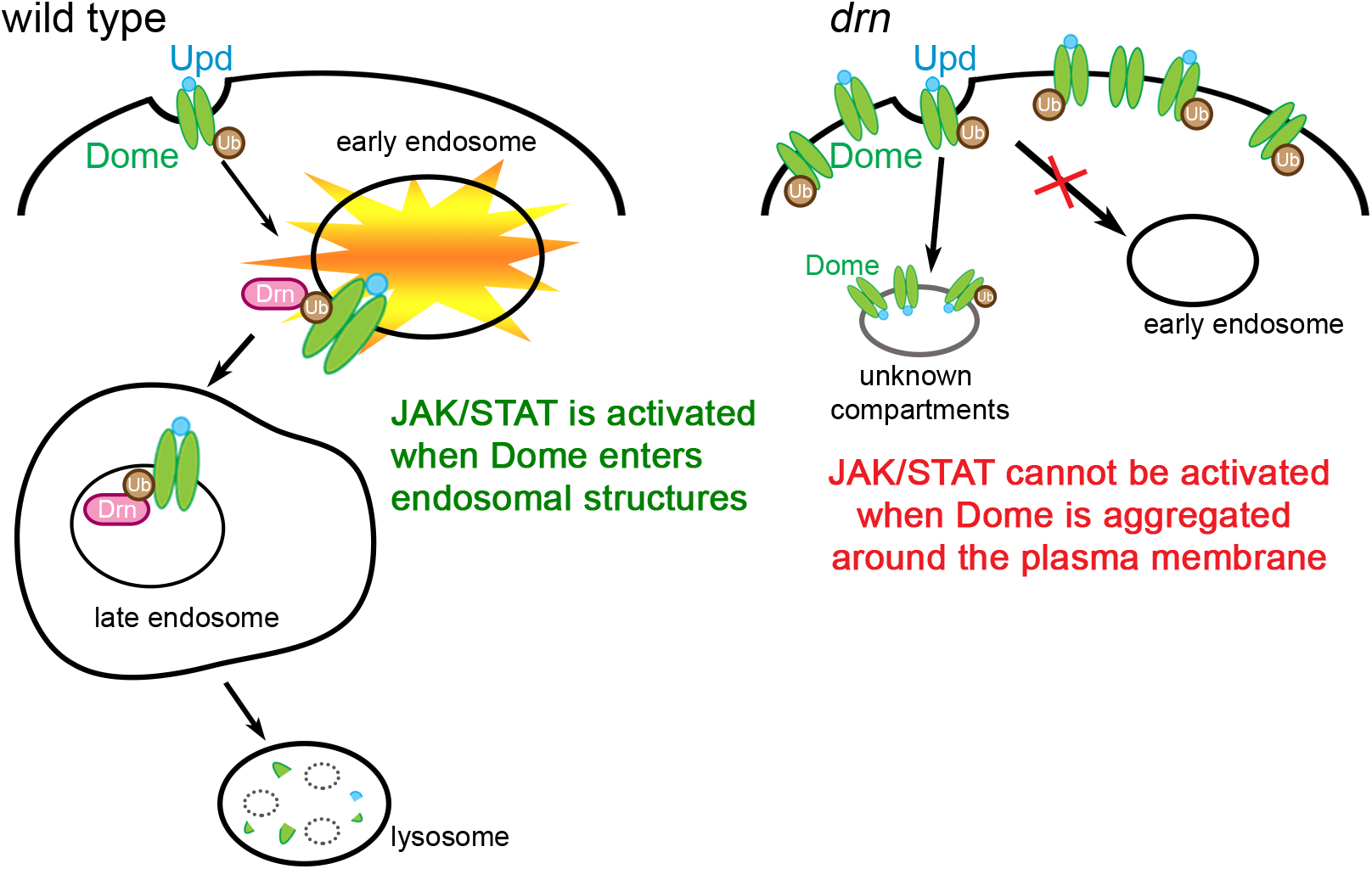
Drn is required for Dome endocytosis, which contributes to the activation of JAK/STAT signaling. A schematic diagram showing the potential role of Drn in the endocytic trafficking of Dome. Ubiquitinated Dome is transported to late endosomes through the endocytic pathway. The direct or indirect interaction of Drn with ubiquitinated Dome may be required for the internalization of Dome, a process that enables the activation of Dome and STAT. Such internalization is also required for Dome’s proper degradation in lysosomes. In contrast, in the absence of Drn, Dome fails to properly enter the endocytic trafficking pathway, which attenuates JAK/STAT signaling activity. Additionally, the failure of endocytosis leads to the abnormal accumulation of Dome.

## DISCUSSION

### Drn contributes to the endocytic trafficking of Dome, which may be coupled with the activation of JAK/STAT signaling

In this study, we demonstrated that *drn*, which encodes a *Drosophila* ortholog of AWP1, plays crucial roles in the LR-asymmetric development of the FG and AMG by positively regulating JAK/STAT signaling. This process requires wild-type *drn* for normal endocytic trafficking of Dome, which is the *Drosophila* JAK/STAT signaling receptor. Whether the internalization and endocytic trafficking of Dome is required to activate JAK/STAT signaling in *Drosophila* has been the subject of much debate (Devergne et al., 2007; Moore et al., 2020; Ren et al., 2015; Vidal et al., 2010). Analyses of mutations affecting Dome endocytosis reveal that Dome must be internalized into early endosomes to activate JAK/STAT signaling (Devergne et al., 2007). In contrast, the RNAi-mediated knockdown of genes required for Dome endocytosis enhanced JAK/STAT signaling activity, demonstrating that Dome endocytosis negatively regulates JAK/STAT signaling (Vidal et al., 2010). This discrepancy can be explained by the multiple ramifications and parallelism of the endocytic pathway, as described in recent studies (Cendrowski et al., 2016); namely, blocking a particular step along the pathway can divert endocytic trafficking into various positive or negative regulatory cascades of cell signaling pathways. The relative contributions of endocytic regulators can also change the course of endocytic trafficking, as is well documented in Notch signaling (Shimizu et al., 2014), and a change in the balance of regulators may change how endocytosis regulates cell signaling pathways according to context. Hence, to comprehensively understand how endocytosis contributes to the activation of Dome, we have to examine the point at which endocytosis is disrupted and the altered course of endocytic trafficking when normal endocytosis fails.

A study using RNAi in cultured *Drosophila* cells identified *drn* as negative regulator of JAK/STAT signaling, which appears opposite to our model (Vidal et al., 2010). However, considering that other genes required for Dome endocytosis were also identified as negative regulators of JAK/STAT signaling in that analysis, it is likely that this discordance regarding the role of Drn in JAK/STAT signaling also results from the discrepancy in contribution of Dome endocytosis to JAK/STAT signaling as discussed above (Cendrowski et al., 2016). Regarding this discrepancy, it was proposed that long-term loss-of-function analyses, including analyses of mutants in which Dome endocytosis is disrupted, may reflect unexpected cell-fate changes induced by altered endocytic pathways and their influence on JAK/STAT signaling (Vidal et al., 2010). However, in this study, we examined three developmental contexts in which JAK/STAT signaling is reduced in *drn* mutants. One of these involves the regulation of *eve* expression, which occurs before cell-fate specification and during a short period of time in early embryogenesis and thus does not fit well with a mechanism involving unexpected cell-fate changes. Therefore, we here propose a model in which Drn is required for some steps of the endocytic trafficking of Dome, which is required for the subsequent activation of JAK/STAT signaling (Fig. 6). Our model is consistent with that proposed by Devergne et al., in which the endocytic trafficking of Dome serves to activate JAK/STAT signaling and also provides a mechanism to regulate it quantitatively (Devergne et al., 2007). However, since it is not clear where and how Drn controls the endocytic trafficking of Dome, we should be cautious in interpreting the coincidence between previous and current results.

### Drn is specifically involved in the endocytic trafficking of Dome

*drn* is the *Drosophila* ortholog of *AWP1*, which binds ubiquitin and modulates the functions of ubiquitinated proteins in mammals and *Xenopus* (Chang et al., 2011; Duan et al., 2000; Fenner et al., 2009; Miyata et al., 2012; Seo et al., 2013). Ubiquitination of the receptors involved in JAK/STAT signaling is important for regulating signaling activities in both mammals and *Drosophila* (Gesbert et al., 2005; Martinez-Moczygemba et al., 2007; Tognon et al., 2014; Wölfler et al., 2009). The sorting of ubiquitinated membrane proteins into the intraluminal vesicles relies on protein complexes in the ESCRT family (Babst et al., 2002a; Babst et al., 2002b; Katzmann et al., 2001). ESCRT-0, I, and II include multiple ubiquitin-binding proteins and interpret ubiquitin as a signal to sort membrane proteins (Clague et al., 2012). In *Drosophila* mutants of the ESCRT-0 complex components *Hrs* and *Stam*, ubiquitinated membrane proteins such as Notch and Dome aggregate at the cell cortex and in intracellular compartments (Jékely and Rørth, 2003; Tognon et al., 2014). We here showed that a loss of Drn caused Dome to accumulate in large clumps that were not labeled by markers for typical intracellular compartments (Fig. 5C-J”). Drn occasionally colocalized with markers of various endocytic compartments, demonstrating that Drn is an endocytic protein (Fig. 4). Hence, we speculated that Drn might play a role in the ubiquitin-dependent internalization or sorting of Dome through Drn’s potential ubiquitin-binding activity (Fig. 6). This idea is consistent with our observation that Drn colocalizes with Dome in some intracellular vesicles in wild-type *Drosophila* (Fig. 5K-K”). Thus, in our model, Dome may be misrouted to endocytic compartments where, in the absence of Drn, it fails to be phosphorylated. Conversely, differences in Dome trafficking routes between wild-type and *drn* mutant embryos should help to identify the endocytic compartment where Dome is activated via phosphorylation. However, it is difficult to delineate incorrect Dome trafficking routes in the *drn* mutant since Dome did not specifically colocalize with typical markers of endocytic compartments under this condition. Our model also predicts that such misrouting consequently prevents Dome’s degradation in lysosomes, leaving it to accumulate as observed in *drn* mutants (Fig. 6).

Our analyses revealed that *drn* mutations induced marked accumulations of Dome but not of Notch or Fz2 (Fig. S5). Notch and Fz2 intracellular distribution in the *drn* mutant appeared similar to wild type. Thus, Drn is not a general component of endosomal protein sorting, but is specific to Dome, though it is not clear how such specificity is achieved.

### Roles of JAK/STAT signaling in LR-asymmetric development of the *Drosophila* gut

We here showed that JAK/STAT signaling activity must be maintained at proper levels for normal LR-asymmetric development of the FG and AMG. We previously observed a similar phenomenon in Wnt or JNK signaling activity (Kuroda et al., 2012; Taniguchi et al., 2007). However, we failed to detect any LR asymmetry in activity or in the distribution of the molecules involved in the JAK/STAT, Wnt, and JNK signaling pathways (Fig. S6) (Kuroda et al., 2012; Taniguchi et al., 2007). Given that these three signaling pathways are required for the LR-asymmetric rearrangement of nuclei in the visceral muscles of the midgut, they may play permissive roles in rearranging these nuclei upon a common cue, as yet unknown, of LR polarity.

AWP1/Drn is highly conserved from *Drosophila* to humans (Fig. 1E). Considering that receptors in the mammalian JAK/STAT pathway are also ubiquitinated, we speculate that the role of AWP1/Drn in JAK/STAT signaling and LR-asymmetric development may be evolutionarily conserved in various organisms (Gesbert et al., 2005; Martinez-Moczygemba et al., 2007; Wölfler et al., 2009).

## MATERIALS AND METHODS

### Fly stocks

We used Canton-S as the wild-type *Drosophila* strain. We generated the *drn*^*1*^ and *drn*^*2*^ mutants in this study. *drn*^*GS12294*^ and *drn*^*GS10567*^ are previously reported GS lines (Toba et al., 1998). *UAS-drn* and *UAS-myc-drn* were generated in this study. *UAS-hop*^*Tuml*^ (Harrison et al., 1995) and *UAS-dome-GFP* have been described. (Ghiglione et al., 2002) *UAS-drnRNAi* (VDRC#103508) was used for RNAi against *drn*. This study used the following GAL4-lines: *da-GAL4* (Wodarz et al., 1995), *48Y-GAL4* (Martin-Bermudo et al., 1997), *24B-GAL4* (Brand and Perrimon, 1993), *hand-GAL4* (Popichenko et al., 2007), *elav-GAL4* (Yao and White, 1994), *NP1522* (Hayashi et al., 2002), *NP5021* (Hayashi et al., 2002), *NP0221* (Hayashi et al., 2002), and *65E04* (Jenett et al., 2012). Lines used to generate homozygotes of *drn*^*2*^ lacking its maternal contribution are *P{ry*^*+t7*.*2*^*=neoFRT}82B ry*^*506*^ (Bloomington #2035) and *w*^*\**^*;P{ry*^*+t7*.*2*^*=neoFRT}82B P{w*^*+mC*^*=ovoD1-18}3R/st*^*1*^*βTub85D*^*D*^*ss*^*1*^*e*^*s*^*/TM3, Sb*^*1*^ (DGRC#106675).

All fly stocks were maintained on a standard *Drosophila* medium at 25 °C unless otherwise stated. Mutant alleles of the second and third chromosome were balanced with appropriate blue-balancers, such as *CyO, P{en1}wg*^*en11*^, *TM6B, AbdA-lacZ*, and *TM3, ftz-lacZ*.

### Generation of *drn*^*1*^ and *drn*^*2*^ mutants

We generated the *drn*-deletion mutants *drn*^*1*^ and *drn*^*2*^ by imprecise excision of P-elements from *GS12689* and *GS10487*, respectively (Toba et al., 1998). Imprecise excision was performed using a standard procedure described previously (Hummel and Klämbt, 2008). The *drn*^*1*^ and *drn*^*2*^ mutations contain deletions from 9,353,341 to 9,355,167 and from 9,353,293 to 9,358,171, respectively (FlyBase2015_03, Dmel Release 6.06).

### Generation homozygotes for *drn* lacking its maternal contribution

We obtained *drn*^*2*^ homozygous embryos lacking the *drn* maternal contribution (*drn*^*2 m/z*^) using standard crosses described previously (Prudencio and Guilgur, 2015). First, *FRT82B* was introduced into the *drn*^*2*^ chromosome by recombination, and flies carrying *FRT82B drn*^*2*^ were selected by Geneticin (Gibco) and genomic PCR using the primers 5’- TCACGCATTCAGAGCTTCGTGTGCCC-3’ and 5’- ATGTTGCTGCGTTTGCTCTGCGTATTCCAC-3’. *FRT82B drn*^*2*^*/TM3b, Sb* females were crossed with *hsFLP/Y; FRT82B ovoD/TM3, Sb* males to obtain *hsFLP/+; FRT82B ovoD/FRT82B drn*^*2*^ females through heat-shock treatment. These females were crossed with *FRT82B drn*^*2*^*/TM3b, Sb* males, and heat shock was performed again to obtain *FRT82B drn*^*2*^ homozygous embryos without the *drn* maternal contribution.

### Generation of UAS*-drn* and UAS*-myc-drn* transgenic flies

To construct *UAS-drn*, a cDNA fragment composed of an entire open reading frame of a *drn* transcript (CG45050-RC) was PCR-amplified using an upper-strand primer containing an *EcoRI* site (5’-CCGGAATTCAGCAGGAAGCAGACGAAACT-3’) and a lower-strand primer containing *HindIII* and *BglII* sites (5’- CCCCAAGCTTAGATCTTCCTTGTTATAGCGCAGCAT-3’). The cDNA clone RE70963 was used as a template (Stapleton et al., 2002). The PCR product was digested with *EcoRI* and *HindIII*, subcloned into the *EcoRI* and *HindIII* sites of pBluescript, and sequenced (Agilent Technologies). The cloned fragment was subcloned into the *EcoRI* and *BglII* sites of the pUAST vector (Brand and Perrimon, 1993).

To construct UAS-*myc-drn*, a DNA fragment composed of an entire open reading frame of a *drn* transcript (CG45050-RC) was PCR-amplified using RE70963 cDNA as a template, an upper-strand primer containing *EcoRI* and *BglII* sites and the *myc*-tag coding sequence (underlined) (5’- CCGGAATTCCAAAATG<u>GAGCAGAAGCTGATCTCGGAGGAGGATCTG</u>AGATCTA TGGAACGTGAATCTAACCC), and a lower-strand primer containing a *XhoI* site (5’- CCGCTCGAGTCAAATCTTTTGAATCTTCT-3’). CG45050-RC has the same open reading frame as that in CG45050-RB, -RD, -RE, -RF, and -RG (Fig. 1C). The PCR product was digested with *EcoRI* and *XhoI* and subcloned into pUAST, and the DNA sequence of the coding region was confirmed. UAS-*drn* and UAS-*myc-drn* constructs were introduced into the *Drosophila* genome using P-element-mediated transformation (Spradling and Rubin, 1982).

### Generation of an anti-Drn antibody

A fragment of *drn* cDNA (RE70963) containing an entire open reading frame of CG45050-RC was amplified by PCR, sequenced, and subcloned into the *BamHI* and *EcoRI* sites of the pGEX-2T vector (GE Healthcare Life Sciences). A GST-Drn fusion protein was produced in Origami B (DE3) cells (Novagen) and purified with a glutathione-Sepharose 4B column. The purified GST-Drn fusion protein was used to immunize rats, and polyclonal antiserum was purified using a standard protocol.

### Antibody staining, *in situ* hybridization, and microscopic analysis

Embryos were immunostained as described previously using the following primary antibodies: mouse anti-Fas3 (1:100, DSHB), chicken anti-β-galactosidase (1:500, Abcam), mouse anti-Pdi (1:200, Stressgen), anti-lectin-PNA (1:500, Vector Laboratories), mouse anti-Sec5 (1:200, 22A2; see Murthy et al., 2003), rabbit anti-GM130 (1:50, Abcam), guinea pig anti-Rab5 (1:3000, gift from Nakamura Akira), rabbit anti-Rab7 (1:5000; see Tanaka and Nakamura, 2008), rabbit anti-Rab11 (1:5000; see Tanaka and Nakamura, 2008), guinea pig anti-Hrs (1:1000; see Lloyd et al., 2002), mouse anti-extra domain of Notch (1:500, DSHB), mouse anti-Frizzled2 (1:20, DSHB), rabbit anti-GFP (1:500, MBL), rabbit anti-RFP (1:500, MBL), rat anti-GFP (1:500, Nacalai Tesque), mouse anti-eve (1:20, DSHB), and rat anti-Drn (1:500). The chitin-binding probe (CBP546) was prepared from a bacterial expression construct using the protocol provided by Yinhua Zhang (New England Biolabs) (Dong et al., 2014). CBP546 (1:50) was added along with secondary antibodies of other primary antibodies for trachea staining. Images were obtained with an LSM880 (Carl Zeiss) microscope and processed with Adobe Photoshop. For *in situ* hybridization, we used standard protocols as described previously (Jiang et al., 1991) and obtained images using an Axiosop2 Plus (Carl Zeiss) microscope.

## ACKNOWLEDGEMENTS

We thank Dr. Norbert Perrimon for providing the *UAS-hop*^*Tuml*^ fly line and Stephane Noselli for providing the *UAS-dome-GFP* fly lines. Fly stocks were obtained from the *Drosophila* Genetic Resource Center at the Kyoto Institute of Technology and the Bloomington *Drosophila* Stock Center at Indiana University. We thank Dr. Hugo Bellen and Dr. Akira Nakamura for providing antibodies. Some antibodies used in this study were obtained from the Developmental Studies Hybridoma Bank (DSHB) at the University of Iowa.

